# The circadian oscillator analysed at the single-transcript level

**DOI:** 10.1101/2020.08.26.268201

**Authors:** Nicholas E. Phillips, Alice Hugues, Jake Yeung, Eric Durandau, Damien Nicolas, Felix Naef

## Abstract

The circadian clock is an endogenous and self-sustained oscillator that anticipates daily environmental cycles and coordinates physiology accordingly. While rhythmic gene expression of circadian genes is well-described in populations of cells, the single-cell mRNA dynamics of multiple core-clock genes remain largely unknown. Here we use single molecule fluorescence in-situ hybridization (smFISH) at multiple time points to measure pairs of core-clock transcripts, *Rev-erbα* (*Nr1d1*), *Cry1* and *Bmal1*, in mouse fibroblasts at single-molecule resolution. The mean mRNA level oscillates over 24 hours for all three genes, but mRNA numbers show considerable spread between cells. While transcript number scales with cell size for all genes, gene-to-gene correlations of mRNA number depends on the gene pair. To account for these features of the data, we develop a probabilistic model for multivariate smFISH mRNA counts that quantifies changes in transcriptional bursting across genes and over circadian time. We identify a mixture model of negative binomials as the preferred model of the mRNA count distributions, which accounts for cell-to-cell heterogeneity, notably in cell size. The paired count data and modelling allows the decomposition of mRNA variability into distinct noise sources, showing that circadian clock time contributes only a small fraction of the total variability in mRNA number between cells. Thus, our results highlight the intrinsic biological challenges in estimating circadian phase from single-cell mRNA counts and suggest that circadian phase in single cells is encoded post-transcriptionally.

## Introduction

In animals, the circadian clock is a 24-hour period oscillator that dynamically regulates central aspects of physiology across all scales of biological organization, from metabolism and locomotor activity down to cellular gene expression and cell signalling [1–3]. Circadian rhythms are entrained by external signals (Zeitgebers) but are present even in single cells; in fact, live-cell imaging of transcriptional reporters [4] and endogenous fusion proteins [5–7] have revealed that circadian oscillations are cell-autonomous and self-sustained, with a period that fluctuates from cycle to cycle by typically 10%. Molecularly, it is thought that the oscillations involve interactions between core circadian clock proteins including BMAL1, CRY1 and REV-ERBα (encoded *by Arntl, Cry1*, and *Nr1d1* genes, respectively), establishing negative feedback loops [8]. Live-cell studies of circadian oscillators in individual cells have thus far remained limited to one gene product at a time, and hence the properties of circadian oscillators in single cells across multiple genes remain largely uncharacterized [4,5].

We thus aimed at characterizing circadian oscillators at the transcript level by using multi-channel mRNA smFISH, which is a sensitive measure of single-cell transcript counts for multiple genes simultaneously [9,10]. While such smFISH measurements provide transcript counts at a single snapshot, the resulting mRNA distributions contain rich information about the underlying dynamical processes. Specifically, the mRNA distributions can be analysed using the telegraph model of transcription [11,12] to estimate the transcriptional bursting kinetics of individual genes using either smFISH [13–17] or even scRNA-seq count data [18–20], though the latter is less sensitive. At steady-state, the distribution predicted by the model, a Beta-Poisson mixture [18,21], can be approximated with a negative binomial (NB) distribution, which is valid when the mRNA half-life is long in relation to the time spent in the active promoter state [13] and which is typical for mammalian genes [22,23]. The NB distribution, which is over-dispersed (having larger variance than expected from a Poisson distribution), uses two informative parameters specifying its shape: the burst size and the burst frequency (normalized by mRNA half-life). Using the telegraph model, the transcriptional parameters for the core clock gene *Bmal1* have been analysed both by smFISH [14] and live imaging of destabilized *Bmal1-Luc* transcriptional reporters [22,23]. These studies showed that the transcriptional burst frequency of *Bmal1* and the clock-output gene *Dbp* oscillates over the circadian clock [14], while the burst size stayed constant.

In addition to gene expression variability caused by biomolecular birth-death processes and transcriptional bursting, both of which contribute intrinsic noise, significant differences in mRNA number may also be caused by extrinsic sources of cell-to-cell variability [24,25], such as cell size or cell-cycle stage [26–28]. Notably, transcriptional burst size has been proposed to correlate with cell volume in both mammalian cells [29] and yeast [30], i.e. transcriptional burst sizes are larger in larger cells. For oscillatory transcripts of the circadian clock, and also transcripts driven by it, variability in the single-cell state of the circadian oscillator (partial synchronisation of the circadian phases) would also further increase transcript count variability in cell populations [31]. The extent to which each of these sources of noise contributes to cell-to-cell heterogeneity in mRNA expression remains an open question for circadian clock genes.

Here, we aimed to quantitatively study how the mRNA distributions of core circadian genes evolve over the circadian cycle while considering multiple sources of intrinsic and extrinsic variability. We used smFISH to simultaneously target pairs of core clock transcripts (*Cry1* and *Nr1d1*, or *Cry1* and *Bmal1*) in confluent NIH3T3 mouse fibroblasts every 4 hours over the circadian clock. We mathematically modelled the mRNA counts using mixtures of distributions to account for intrinsic transcriptional fluctuations, cell-to-cell variability and time-varying (periodic) parameters. We investigated several models of increasing complexity and, using a Bayesian model selection approach, we found that the preferred model favours inclusion of measured cellular area as an explanatory variable, with gene-specific scaling of mRNA counts with cell size. From the preferred model we could decompose the sources of measured variation in core clock transcript number, showing that only a small percentage (∼5%) is caused through circadian time. Due to the strong contribution of transcriptional bursting in individual genes, our results suggest that circadian phase in individual cells may be specified by mRNA numbers of many genes together, or by other molecular states such as protein abundance or protein activity levels.

## Results

### Time-resolved mRNA count distributions of core clock genes in single cells

To characterise the circadian oscillator at the single-transcript level, we performed smFISH in synchronised, confluent (non-dividing) NIH3T3 mouse fibroblasts and measured transcript numbers every 4 hours for 24 hours (7 time points) (Fig. 1A). The cells were synchronised using dexamethasone (Dex), and sampling started 17 hours after the treatment to avoid the initial transient response. We performed multi-channel imaging with fluorescent probes targeting exons of *Bmal1* and *Cry1* or *Nr1d1* and *Cry1* in the same cells, and we measured the number of transcripts per cell for each gene in approximately 450 single cells per time point (Methods). The mean number of *Bmal1, Cry1*, and *Nr1d1* transcripts per cell oscillated along the circadian cycle and ranged from 10-35 molecules (Fig. 1B), where the average copy number of *Bmal1* was in the same range as previous measurements in the same cell line [14]. To estimate the phase and amplitude of each gene we fitted a two-harmonic cosinor model (Methods). The oscillations of *Cry1* and *Bmal1* are approximately in anti-phase with *Nr1d1* positioned in-between, which is consistent with known phase relationships of core clock transcripts in 3T3 cells in bulk measurements [32,33] (Fig. 1C). The fold change, defined as the ratio of the peak to the trough, was 1.50 for *Cry1*, 1.57 for *Bmal1* and 2.05 for *Nr1d1* (Fig. 1C). Even though the cells were synchronised with Dex, we still expect cell-to-cell differences in the phase resulting from incomplete synchronisation [31]. We therefore entrained cells with temperatures cycles, known to be efficient method of synchronisation, and obtained a fold change of 2.1, which was slightly higher than with Dex synchronisation and similar to previous reports [34] (Supp figure 1).

**Figure 1:**
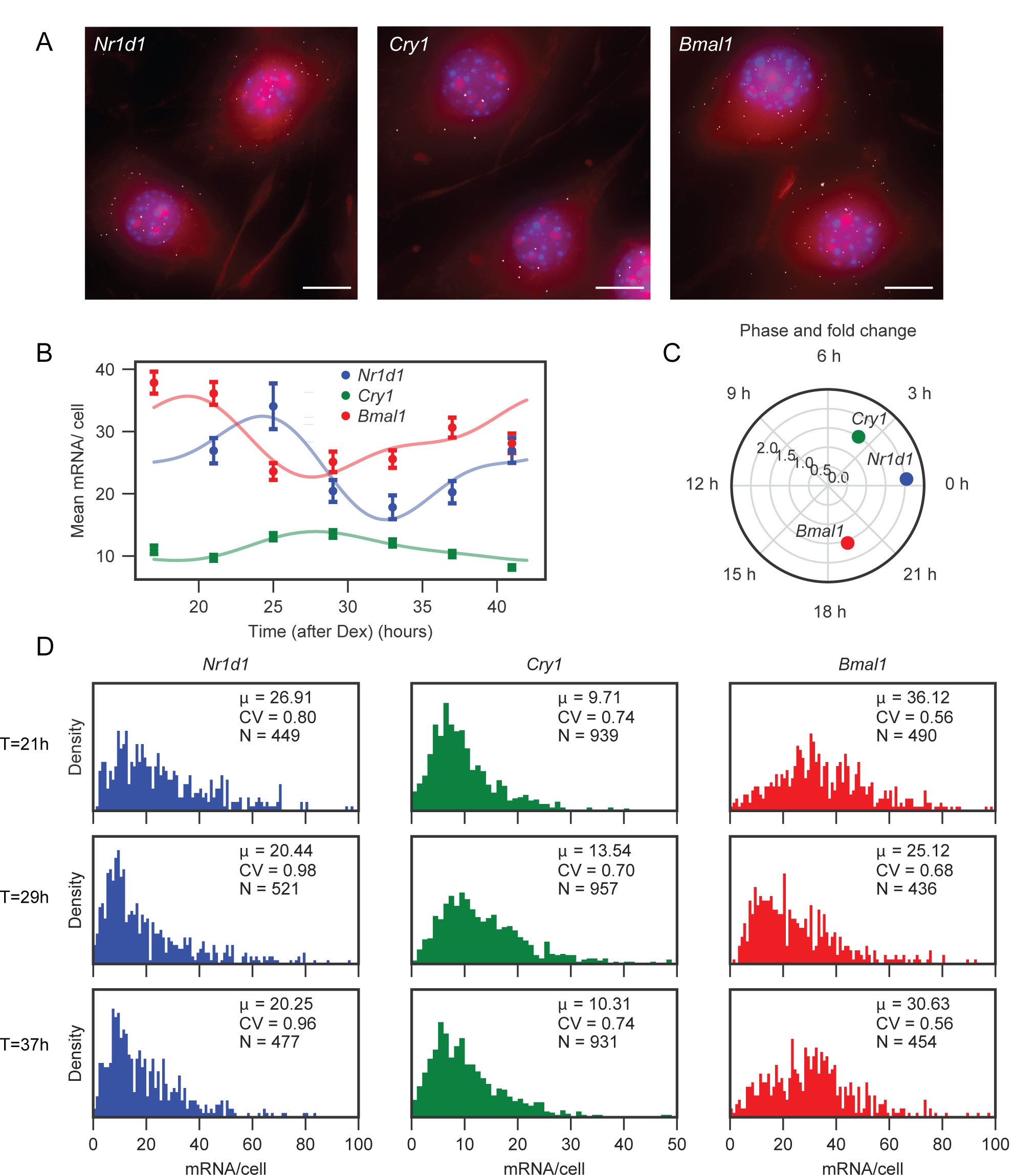
Single-Molecule RNA Fluorescence In Situ Hybridisation (smRNA FISH) captures transcript distributions of core clock genes in mouse fibroblasts at multiple time points. (A) smFISH targeting *Bmal1, Cry1* and *Nr1d1* in wild-type NIH 3T3 cells at 17h, 25h and 17h after synchronisation with Dex, respectively. Each image corresponds to a maximum projection of the z-stack. Nuclei are stained with DAPI (blue) and cytoplasms with HCS CellMask (red). Each fluorescent dot (white) corresponds to a single transcript. Scale bar: 20 μm. (B) Dots, the average number of *Bmal1, Cry1*, and *Nr1d1* transcripts per cell as a function of time after treatment with Dex. Error bars represent the standard error on the mean. The solid lines represent fits using a two-harmonic cosinor model (Equation (1), Methods) for each gene individually. (C) The inferred peak phase (angular component of graph) and max-to-min fold change (radial component) from the fit of Equation (1). (D) Distributions of *Nr1d1, Cry1* and *Bmal1* transcripts at 21h, 29h and 37h after synchronisation with Dex. *μ* represents the mean of the distribution, CV represent the coefficient of variation (standard deviation / mean), and N is the number of cells at the given time point.

While the mean mRNA level changed over time for all genes, there was substantial variability in mRNA numbers between cells, leading to significant overlap of the transcript distribution between different time points (Fig. 1D). The measured coefficients of variation (CV = standard deviation / mean) ranged from 0.5-1. For comparison, the expected CVs would be 0.17-0.32 if a gene had an average mRNA level between 10-35 molecules and if mRNA dynamics followed a Poisson process (i.e. assumes constant mRNA production and degradation rates and cells share the same kinetic parameters). Thus, the observed variability implies additional contributions, possibly from transcriptional bursting or extrinsic sources [26]. The mathematical modelling introduced below serves to identify these contributions.

### Core clock transcript numbers scale with cell size

Given that previous studies identified cell size as a source of extrinsic variability in transcript counts [26,27,29,35], we quantified cell sizes and found a positive correlation with mRNA number for all genes (Supp figure 2). In fact, we found a linear relationship between the log area and the log mean mRNA number, indicating a power-law scaling (Fig. 2A). The relationship was sub-linear, with exponents ranging from 0.42 for *Nr1d1* to 0.70 for *Bmal1*. If mRNA count were proportional to volume and if the cell had a geometry such as a sphere or cube, we would expect a super-linear scaling between mRNA count and area. The observed sub-linear scaling thus likely reflects the “fried egg” morphology of adherent cells, where increases to the area of a relatively flat cytoplasm have a proportionally small effect on the total volume due to the nucleus.

**Figure 2:**
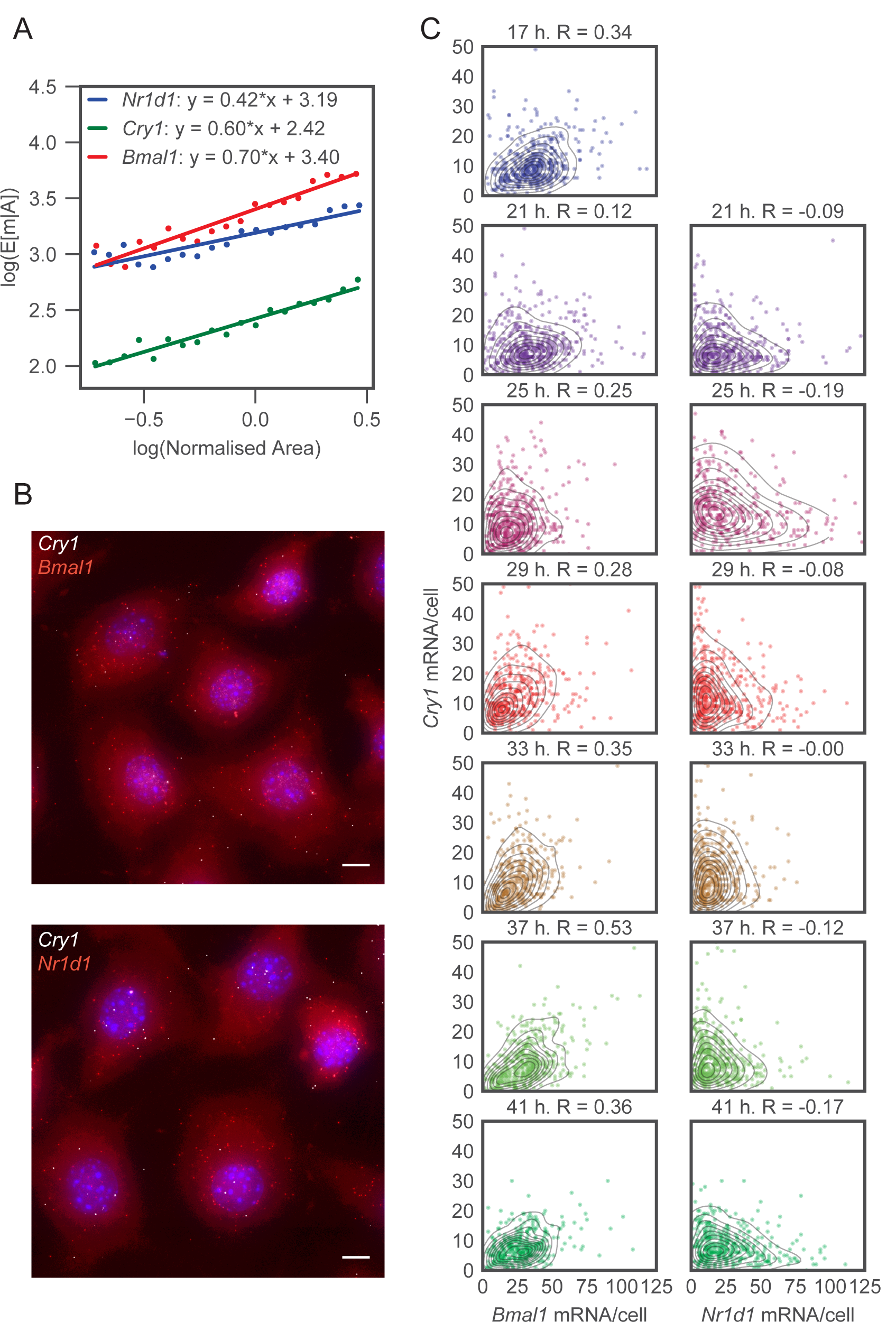
Core circadian transcript distributions show area dependence and distinct gene-to-gene correlation structures. (A) Log conditional mean mRNA count for a given area as a function of the log area. Each cell is sorted by area by placing it in one of 20 bins that range from the 5^th^ to the 95^th^ percentile. The conditional mean given the area E[m|A] is then the mean mRNA count of each bin. The area is normalised such that the average area is equal to one. Solid lines represent a least-squares linear fit to the data. (B) Top image: Dual-channel smFISH targeting *Bmal1* and *Cry1* simultaneously, taken at 21h after Dex synchronisation. Blue, nuclei stained with DAPI; white dots, *Cry1* transcripts; magenta, *Bmal1* transcripts. Bottom image: Dual-channel smFISH targeting *Nr1d1* and *Cry1* simultaneously, taken at 21h after Dex synchronisation. Blue, nuclei stained with DAPI; white dots, *Cry1* transcripts; magenta, *Nr1d1* transcripts. Scale bar: 20 μm. (C) Bivariate distributions of mRNA counts per cell for dual-channel smFISH targeting either *Bmal1* and *Cry1* or *Nr1d1* and *Cry1*. Each dot corresponds to a single cell, and the contours represent KDE estimates of the density. Time represents the number of hours after Dex synchronisation, and R represent the Pearson correlation coefficient.

### Two-gene mRNA count distributions reveal gene-pair specific correlations

We next exploited the ability of smFISH to measure multiple transcripts simultaneously to explore the joint relationship between transcript numbers of different clock genes. Our dual-channel imaging allows either *Bmal1*/*Cry1* or *Nr1d1*/*Cry1* to be measured in the same cells (Fig. 2B). The bivariate relationships between the gene pairs show that *Bmal1*/*Cry1* are positively correlated at each time point (R from 0.12 to 0.53), whereas *Nr1d1*/*Cry1* show negative correlations (R from 0.0 to -0.19) (Fig. 2C). Since all genes are positively correlated with area (Fig. 2A), the negative correlation between *Cry1* and *Nr1d1* could be caused by a spread in the phases between cells [36], or regulatory interactions (e.g. NR1D1 protein represses *Cry1* transcription), which can cause different steady-state correlations depending on whether feedback is negative or positive [11]. Below, we formulate these hypotheses as simplified, effective mathematical models to quantify the compatibility of our data with both scenarios.

### Modelling periodic mRNA count distributions in heterogeneous cell populations as mixtures of negative binomials

We next developed a model with the aim of finding a compact mathematical representation of the circadian clock in the space of multi-dimensional transcript counts. In other words, we sought to describe how the multi-variate probability distribution of mRNA counts varies as a function of circadian time. As the above exploratory analysis shows, there are systematic effects (e.g. cell features) coupled with time-varying parameters (the circadian oscillator), and the variance of the mRNA distributions is larger than expected with a Poisson process. To keep the model manageable and interpretable we made a number of simplifying assumptions. First, since time scales associated with transcriptional bursting as well as mRNA half-lives of clock genes are short compared to the 24-h period, we modelled the system in a quasi-steady state, which means that at each point the mRNA count distribution in the population is approximated with a slowly time-varying stationary distribution. Second, based on reports that transcriptional bursts are typically short relative to the mRNA half-life [22,23], we used a common approximation of the full telegraph model which takes the form of an NB distribution, with the burst size and burst frequency as the two effective parameters [13]. We also proposed additional model features in four different but related models of increasing complexity (Table 1 and Fig. 3A). We then selected the optimal model from the candidates using Bayesian model selection.

**Table 1:** The features (F1-F5) included in each of the four models (M1-M4) considered.

|  | M1 | M2 | M3 | M4 |
| --- | --- | --- | --- | --- |
| F1: Time-dependent burst frequency | X | X | X | X |
| F2: Constant burst size | X |  |  |  |
| F3: Cell-size dependent burst size |  | X | X | X |
| F4: Cell-specific circadian phase |  |  | X |  |
| F5: Cell-specific burst size |  |  |  | X |

**Figure 3:**
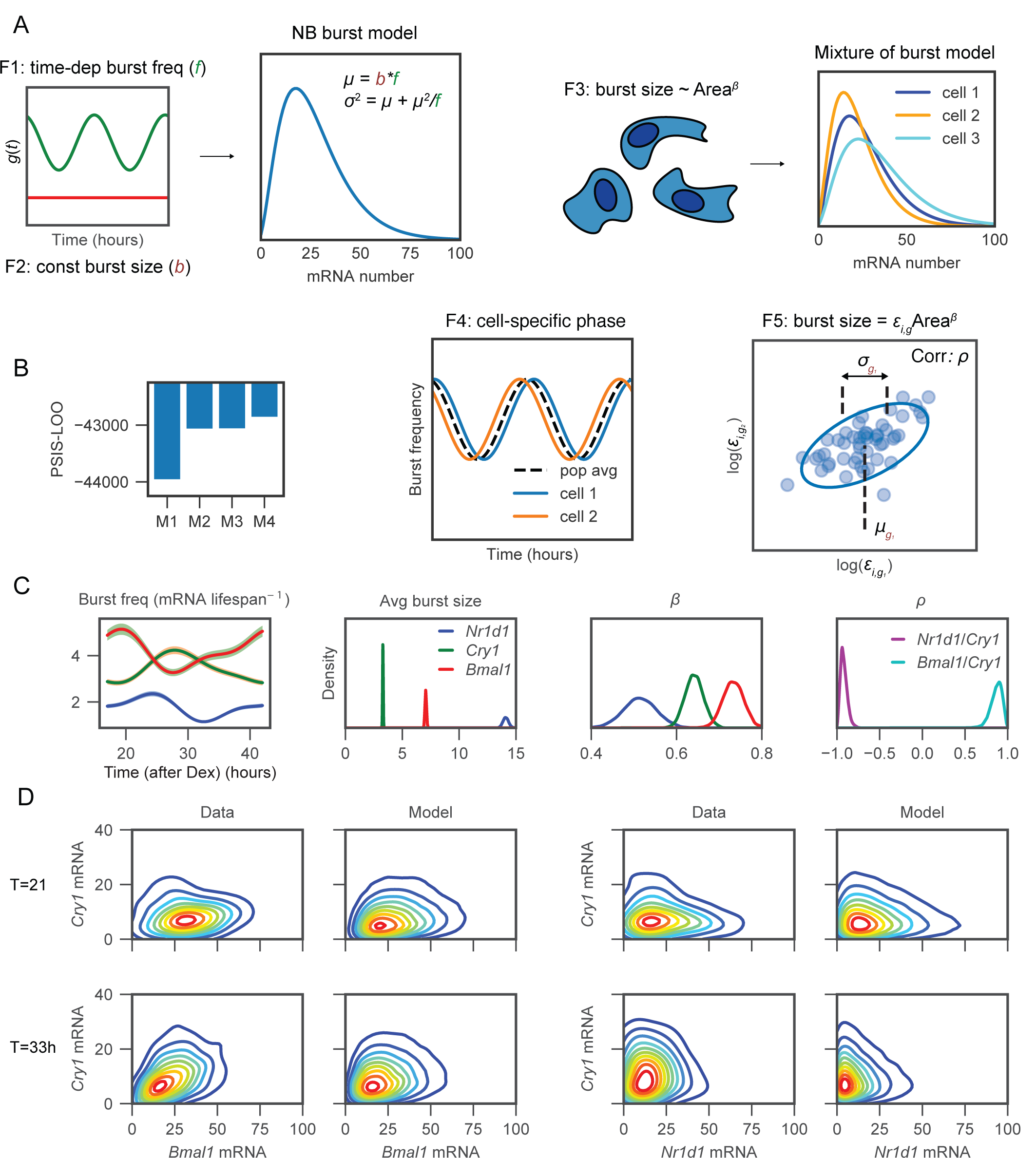
A mathematical model that includes area dependence and gene-specific correlation in burst size captures the observed time-dependent bivariate mRNA distributions. (A) Schematic of the four models considered. (B) PSIS-LOO calculated for each of the four models using the whole dataset of both gene-pairs across all time points. (C) Posterior parameter estimates. Burst frequency is plotted as a function of time, where the solid line represents the posterior mean and the shaded area represents the 90% confidence interval. Posterior probability densities are shown for the average burst size, *β* (which controls the dependence between burst size and cell area) and *ρ* (representing the correlation in burst size between genes (in log space)), where *β* is inferred from model M2 and the average burst size and *ρ* are calculated from model M4. (D) omparison of the probability density of the data (kernel density estimates) with the model (using model M4). To estimate the probability from the model, the data set was simulated 15 times using the posterior mean parameter values combined with the measured cell areas.

The first and simplest model (M1) assumes that the transcriptional bursting parameters controlling the shape of the NB distribution are modulated in a deterministic way by the clock, with no further sources of cell-to-cell heterogeneity. Moreover, we incorporate previous findings regarding transcriptional bursting kinetics of clock genes to further reduce the complexity of the model. Namely, previous results using the same cell line have shown that burst frequency of clock genes is modulated by the circadian clock while the burst size remains constant [14]. In M1, we therefore used an NB distribution with an oscillatory burst frequency but otherwise constant parameters (features F1 and F2, Table 1 and Fig. 3A).

The second model explicitly incorporates cell size as a source of extrinsic variability. Previous work in mammalian cells showed that burst size scales with cell volume [29]. In model M2, we allow the burst size to scale with cell size, and the full mRNA distribution across the population at each time point consequently becomes a mixture of NB distributions (feature F3, Fig. 3A). Since we measured cellular area instead of volume, we set the dependence between burst size and cell area using an additional, gene-specific exponentβ, which is also supported by the linear relationship between log area and the log mean mRNA observed in Fig. 2A.

To compare models M1 and M2, we first inferred parameters using the data for both *Bmal1*/*Cry1* and *Nr1d1*/*Cry1* pairs simultaneously using Hamiltonian Monte Carlo sampling within the STAN probabilistic programming language [37]. We quantified the performance of each model by estimating the out-of-sample predictive accuracy, which rewards a good fit of the model to data while penalising model complexity. We used approximate leave-one-out cross-validation with Pareto-smoothed importance sampling (PSIS-LOO) to estimate the pointwise out-of-sample prediction accuracy (as described in [38]), where models with the highest PSIS-LOO score have the best performance. The PSIS-LOO calculated from both models showed a clear preference for model M2 compared to M1 (Fig. 3B), and we hence built subsequent models using cell-size dependent burst sizes. The dependence on size differed between the three genes (parameter β, Fig. 3C), with *Nr1d1* having the weakest and *Bmal1* the strongest dependency, suggesting that the scaling between mRNA number and cell size for the measured clock transcripts can be gene-specific.

To capture additional features observed in the data, notably the positive and negative correlations in Fig. 2C, we also allowed cells to be imperfectly synchronised (model M3). A spread of circadian phases in a cell population could distort the correlation in mRNA counts between genes, as two genes that are on average in-phase/antiphase but where phases vary between cells could generate positive/negative correlations. To model potentially imperfect synchronisation we introduced cell-specific phases that are distributed around a population average (feature F4, Fig. 3A). To ensure the cell-specific phases do not become too spread we used a von Mises prior with *k* = 2, which approximately matches the phase spread observed in live microscopy for mammalian fibroblasts [31]. To simplify the analysis we fixed parameters previously contained in model M2 to their posterior mean values, and in M3 we keep the waveform shape and phase for the bursting frequency fixed (from M2), but allow the burst frequency amplitude to vary. Fitting Model M3 is practically more difficult and involves many local minima. We therefore performed inference multiple (eight) times and chose the posterior chain with the highest average likelihood. Compared to model M1 the amplitude of oscillations in burst frequency increases by approximately 8%, indicating that the underlying amplitude in single cells is greater than observed at the mean level due to partial synchronisation (Supp figure 3); however, the improvement was minor and the increased complexity of M3 over M2 was not supported according to the PSIS-LOO criterion.

Finally, we considered a further refinement (model M4) that introduces variable burst size from cell to cell as an alternative mechanism of inducing correlation between genes. While in model M2 the burst size was assumed to be directly proportional to the cell area, in model M4 we included an additional cell-specific random variable ε_*i,g*_ to modify the burst size in each cell for each gene (feature F5, Fig. 3A). Model M4 is a hierarchical model whereby the distribution of cell-specific parameters is also learnt during inference, and we assumed that the ε_*i,g*_ is bivariate log-normally distributed between pairs of genes. This log-normal distribution is parameterised with μ_*g*_, σ_*g*_ and ρ, which represent the mean, variance and correlation of ε_*i,g*_ between genes (in log space). Practically, this means that when bursts are large for one gene, they can be large (correlated) or small (anti-correlated) for another gene, and, while the precise mechanism is left unspecified, it can be interpreted as a signature of regulatory interaction (Discussion). When all parameters are free, we noticed that the burst frequency becomes unrealistically high due to a tendency to overfit to individual cells, and we therefore lock the burst frequency values to the posterior mean values from model M2. The PSIS-LOO scores overall favoured model M4 (Fig. 3B), and the predicted joint probability density shows good similarity to the observed data (Fig. 3D) (all time points shown in Supp figure 4). The inferred average burst frequency was highest for *Bmal1* and lowest for *Nr1d1*, while *Nr1d1* showed the highest and *Cry1* the lowest average burst size (Fig. 3C). The burst frequencies (normalised by mRNA half-life) were in the upper range when compared to the transcriptome-wide estimates using single-cell RNA-seq, while the burst sizes spanned the observed range [19]. The correlation parameter ρ of the random variable ε_*i,g*_ was positive for *Cry1*/*Bmal1* and negative for *Cry1*/*Nr1d1* (Fig. 3C), consistent with the observed correlations in the data (Fig. 2C). Overall, we identified a preferred model (M4) that incorporated transcriptional bursting, cell size and circadian time variation, and which was able to capture the measured bivariate smFISH densities.

### Circadian oscillations at the transcript level are blurred by other sources of single-cell heterogeneity

Having selected a preferred model to describe the data, we next analysed the model to decompose the variance into distinct sources. In model M4 there are four sources of transcript count variability across populations of individual cells: temporal control by the circadian clock, intrinsic noise due to the mRNA production-decay process, extrinsic noise from variable cell size, and fluctuations in burst size (ε_*i,g*_, termed “other extrinsic”). Intrinsic noise can be further partitioned into a Poisson component that arises due to the discreteness of mRNA counts (and would be present with constant, constitutive expression) and a second component caused by transcriptional bursting. To decompose the variance, we use the law of total variance (Methods).

For all three genes we found that the variance was dominated by intrinsic noise (Fig 4A), with bursting having a significantly larger contribution than the Poisson component. The extrinsic sources of noise in our model, which include area and variable burst size, had a weaker contribution for all three clock genes. Previous studies have shown that the relative proportions of intrinsic versus extrinsic noise is both condition-(cell types, cell states) and gene-specific, and regression models that use cellular features as explanatory variables can account for between 10-80% of the variance, depending on the gene [26,27]. In the cells used here, the strong influence of transcriptional bursting and dominance of intrinsic noise is consistent with live imaging of a *Bmal1* transcriptional reporter in the same cell line under similar growth conditions, where intrinsic noise was estimated to be 4-times larger than extrinsic noise [23]. As we are only able to quantify the role of extrinsic sources that are included in the model, it is nevertheless possible that inclusion of additional sources such as the cell cycle and phenotypic heterogeneity would increase the proportion of extrinsic noise. Nonetheless, as shown previously [14], cells were maintained in conditions that minimised cell-cycle progression and NIH3T3 fibroblasts are phenotypically homogenous, which controls at least for those major sources of extrinsic noise [27,39].

**Figure 4:**
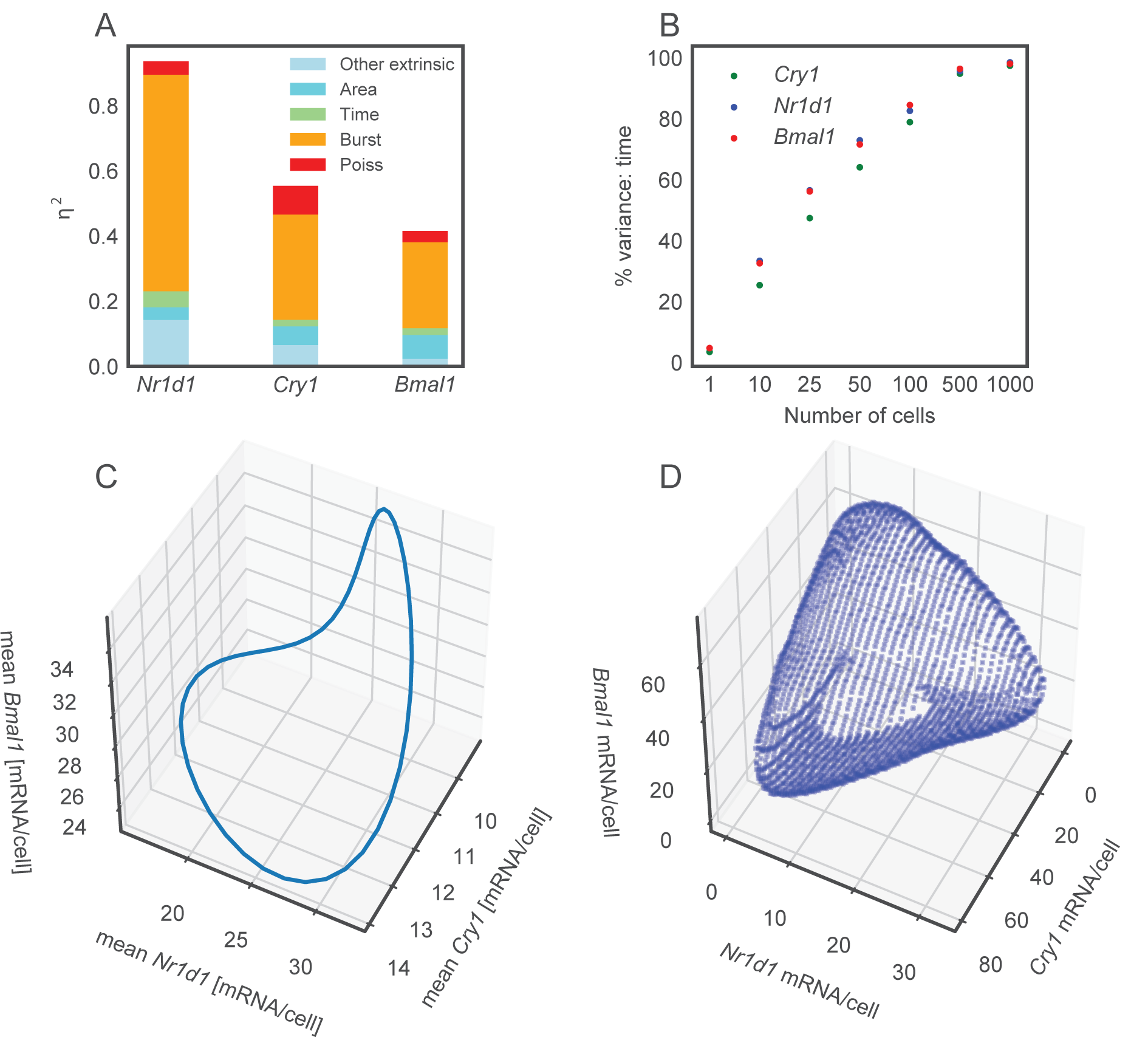
Decomposing the sources of noise shows time has a small contribution to overall mRNA count variance. (A) Noise decomposition of model M4, estimated with 100 simulations of the data set using the posterior mean parameter values combined with the measured cell areas. The variable *η*^2^ represents the variance/mean^2^. (B) Simulations from model M4 to show the percentage of variance attributed to time as a function of the number of cells in each sample. (C) Simulation of the 3-dimensional mean mRNA trajectory of *Bmal1, Nr1d1* and *Cry1*. Each point on the cycle represents a different circadian time. (D) Simulation of the 3-dimensional probability distribution of *Bmal1, Nr1d1* and *Cry1*, averaged over time. The surface represents the area of highest probability density that integrates to 80% of the total probability.

Somewhat surprisingly, the contribution of the 24-h cycle to mRNA variance was low across all genes, consistent with the large overlap in mRNA distributions at different time points (Fig. 1D). This may be unexpected given that oscillations in mRNA levels are routinely detected in both bulk RNA sequencing in tissues [40] as well as cell culture [33,41]. To understand if our results are compatible with circadian rhythms found in bulk cell line RNA sequencing, we calculated the percentage of variance due to time when increasingly large pools of simulated cells are considered (Fig 4B). Starting from a small percentage with one cell (as previously shown in Fig. 4A), the percentage of variance explained by time increases to almost 100% with 1000 cells, which is caused by the increased averaging-out of other sources of single-cell noise as cell numbers become larger. Our results on single cells are therefore compatible with RNA sequencing of bulk populations of cells.

While our experiments and inference were performed on pairs of genes, the model allows us to simulate the count distributions for all three genes simultaneously. As the correlation of ε_*i,g*_ between *Nr1d1* and *Bmal1* is not inferred directly in our model, we use the *Cry1*/*Bmal1* and *Cry1*/*Nr1d1* correlations together with the minimal assumption of conditional independence of *Bmal1* and *Nr1d1* given *Cry1* (Methods). The three-dimensional simulations help visualise the differences between the average and the single-cell measurement of circadian clock mRNA expression. The mean mRNA level shows a clear periodic structure (Fig. 4C), where each circadian time represents a different point on the cycle. In a population of unsynchronised single cells, one might expect noise to blur this structure. It transpires that due to the large contribution of transcriptional burst noise and the weak contribution of time to mRNA variance, the single-cell joint three-dimensional probability distribution is significantly blurred with a diffuse and amorphous structure (Fig. 4D). In summary, probabilistic modelling of the 3-dimensional core clock system shows a 24-hour cycle in the mean level, but this well-defined periodic shape is blurry when the full probability density of mRNA counts is considered.

## Discussion

Single-cell measurement of gene expression with smFISH goes beyond bulk mRNA measurement as it allows for the full characterisation of transcript distributions in populations of cells. Until now, the temporal evolution of circadian mRNA distributions through the clock remained uncharacterized. Here we performed a bivariate measurement of mRNA number across two pairs of genes every 4 hours over the circadian clock. There were several notable features of our data including: an average mRNA expression level that oscillated over the circadian clock, a coefficient of variation for each gene that largely exceeded that from a Poisson process, a transcript number that correlated with cell area and a correlation between genes that depended on the pair considered. To capture these phenomena in a unified quantitative framework, we proposed several probabilistic, generative models of our data and used model selection to balance goodness of fit and model complexity to identify the optimal model.

While we have developed an analysis approach for the circadian clock i.e. a temporal system with a periodic structure, we anticipate that the modelling of smFISH developed in this paper, in particular the ability to include and distinguish intrinsic and extrinsic noise sources, will be relevant to other biological systems. Our starting point was to model the number of mRNA molecules in each cell with a time-varying negative binomial distribution, and by subsequently incorporating cell area the resulting distribution of mRNA counts across the entire population of cells became a mixture of NBs. While previous works have modelled extrinsic noise using cell-specific parameters [42–44], we here specifically used mixtures of NB distributions to model smFISH data. While cell area affected both genes in the same cell (and hence always induces positive correlation), we also introduced cell-specific latent variables (either cell phase or burst size) that can alter the bivariate probability distributions more flexibly. Given the ease with which these types of models can now be implemented in probabilistic programming languages such as STAN, we anticipate that the use of such latent variable models will be instrumental in describing and quantifying the underlying (but unobserved) biology that influences gene-gene relationships in multi-variate single-cell measurements of gene expression.

The considered models for the circadian smFISH data (Table 1) consisted of different mixtures of negative binomial distributions with a burst frequency that changes over time. The final preferred model had a hierarchical structure and included cell-specific burst sizes that were connected at the population level using a bivariate log-normal distribution, which permitted covariance between different genes. The exact mechanistic interpretation is not specified but could possibly arise from regulatory interactions, given that these three clock genes are intertwined in a web of feedback loops. For example, BMAL1 activates *Nr1d1* transcription, CRY1 is known to repress *Nr1d1* transcription by inhibiting BMAL1 activity while NR1D1 directly represses *Cry1* transcription, and hence the observed pair-dependent correlations may be an outcome of this network [8]. While computational methods have recently been developed to exploit cell-to-cell heterogeneity to infer gene regulatory topologies [45–47], building explicit mechanistic models of transcription factor networks remains challenging without additional information on protein levels and activities.

Using the preferred model, we decomposed the variance of the mRNA distributions into distinct components and found transcriptional bursting to be the largest contributor. With further measurements of cellular parameters (e.g. cell volume, mitochondria, cell microenvironment, cell cycle stage), one would expect the estimated contribution of extrinsic noise to grow. However, for the cell type used in our study, cell proliferation was minimised, reducing at least one major source of extrinsic noise [14], and live-cell imaging of *Bmal1* transcriptional reporter indicated that intrinsic noise was dominant [23]. Temporal changes contributed only a small fraction to the total variability in mRNA counts, which is also reflected in the relatively low oscillatory amplitudes we found (Fig. 1B). Though cells that were subjected to temperature entrainment did not yield significantly larger amplitudes (Supp figure 1), one potential cause we have explored in our modelling (M3) is partial synchrony of our cells. The outcome was that partial synchrony was not a likely explanation of the low amplitudes, though it could be that we have not found the true minimum of M3 due to too many local minima. Nevertheless, with CVs between 0.5 and 1, which is typical for mammalian genes expressed on the order of 10 molecules [23,26], even an increased amplitude of 4-fold would represent only ∼38% (CV = 0.5) or ∼12% (CV = 1) of the total variance. In other terms, unless the amplitude fold change is radically increased or the CV decreased, time is unlikely to have a dominant contribution to total variance.

In principle, the possibility of using transcript counts to estimate single-cell circadian phase is attractive and would have important applications in single-cell genomics; however, the fact that time variability is dominated by other sources of variation highlights significant challenges. A possible solution is to leverage mRNA numbers for a large number of genes, for example using single-cell RNA-seq, as successfully done to estimate cell-cycle phase [48–50]. In comparison, analogous methods to estimate circadian phase in single cells currently remain unexplored, although we note though that cell-cycle regulated transcripts are expressed at higher levels compared to core-clock transcription factors.

The fact that circadian time captures only a low contribution to total mRNA variability may be surprising given that clear circadian oscillations of a single-cell reporter (*Rev-Erbα*-YFP) are observed in the same NIH 3T3 cells [4,51] and in human U2OS cells [52]. It should be noted that the reporter used in these cells is probably present in many copies, since the cells were selected for high signals following standard transfection. The multiple promoters could thus resultantly cause an averaging effect, similar to shown in Fig. 4B. Nonetheless, 24-hour rhythms are seen in primary fibroblasts dissociated from *mPer2*-luciferase protein fusion knock-in mice, which shows that robust oscillations are possible at the protein level even when transcribed from diploid alleles. One possible explanation is that there is noise reduction at the protein level. Indeed, using a *Bmal1* transcriptional reporter it was shown that the relative amplitude of oscillations at the luminescent reporter protein level was greater than that observed at the mRNA level [14], which could be caused by regulation of translation or protein degradation over the circadian clock (as known for core clock genes) [22,53]. Mass spectrometry has demonstrated that many mRNAs with flat mRNA profiles over the circadian clock exhibit oscillatory protein expression [54–56], which could be due to the clock regulated translation [57] or degradation. Protein degradation via autophagy can be circadian in mouse liver [58], and rhythmically regulated protein half-lives have been shown to be responsible for phase differences observed between mRNA and protein levels [59,60].

Another important aspect of this question is how circadian phase is encoded at the single-cell level, or, in other words, what are the state variables of the clock. For instance, most of the core clock protein functions seem to converge on regulating the rhythmic activity of the CLOCK-BMAL1 transcription factor complex, thus integrating multiple layers of regulation such as protein stability and dimerization, nuclear import, binding of co-factors and repressors, and phosphorylation of DNA binding domains [61]. Our data suggest that endogenous gene readouts of this activity at the transcript count level are too noisy to faithfully encode single-cell phase, which is however likely defined as a systems property involving activities of protein complexes that are subject to regulation at the post-transcriptional and post-translational levels [61–63].

In sum, our results show that mRNA count distributions of the core clock genes *Bmal1, Nr1d1* and *Cry1* are subject to significant variability such that the circadian limit cycle attractor is not sharply defined at the transcript count level, and future studies should aim to ascertain how single cells are successfully able to generate robust, high-quality circadian oscillations that have been observed in live-cell imaging.

## Methods

### Cell lines and culture condition

For maintenance, wild-type mouse NIH3T3 fibroblasts were cultured in Dulbecco’s Modified Eagle Medium (DMEM, Gibco) complemented with 10% Foetal Bovine Serum (FBS, Sigma) and 1% PSG antibiotics (Gibco). Cells were grown at 37°C, 5% CO2 and 100% humidity and passaged every two to three days, until the day of the experiment. Circadian clock entrainment by temperature cycles was performed using a Memmert INCO153 incubator controlled by the Memmert Celsius 10.0 software. We entrained the clock using temperature cycles ranging from 35.5°C to 38.5°C, as previously described [34] and for at least 10 days prior beginning of the experiment. To maintain synchronisation after passaging, we applied the splitting procedure always at the same temperature cycle position (temp=37°C), when the clock was at its trough. To enhance the synchronisation when splitting, we applied a serum shock by recovering trypsinised cell into DMEM without serum prior inoculating a new dish containing serum supplemented DMEM [64].

### Single-Molecule RNA Fluorescence In Situ Hybridisation (smRNA FISH)

Cells were plated on 6 well-plates containing 18 mm round cover glasses (Fisher Scientific) treated with a solution of 25μg.mL^-1^ Fibronectin (Sigma-Aldrich) diluted in 1X PBS for 30 min at room temperature. Wells were seeded with 0.2× 10^6^ cells then cultured in serum-free DMEM (Gibco), supplemented with 1% PSG antibiotics, to prevent cell division. 10 hours later, cells were treated with 100nM dexamethasone (Sigma-Aldrich) for 30 mins and followed by medium change. For the experiment where the circadian clock was synchronised using temperature cycles, the same procedure was applied except entrained cells were not treated with dexamethasone. The smRNA FISH protocol was largely adapted from the Stellaris RNA FISH protocol for adherent cells, which describes the method used by [10]. Stellaris exonic probes coupled with Quasar570 (548/566nm, Red, BMAL1^Red^, NR1D1^Red^) or Quasar670 (647/670nm, FarRed, BMAL1^FarRed^, CRY1^FarRed^) dyes has been hybridised to targeted mRNA. The hybridisation was performed in 50μL of Stellaris hybridisation buffer complemented with 250μg.mL^-1^ Yeast tRNA (Ambion) and 5mM Ribonucleoside Vanadyl Complex (New England Biolabs). Cells and nuclei were then co-strained with 0.4μg.mL^-1^ green HCS CellMask (Invitrogen) and 0.7μg.mL^-1^ DAPI (Thermofisher), respectively. HCS CellMask and DAPI were diluted in the Stellaris wash buffer. Cover glasses were finally mounted onto microscopy slides in ProLong™ Gold Antifade Mountant (Thermofisher). A total of 3 slides were imaged for each time point.

### Imaging

Cells were imaged at the EPFL imaging facility (BIOP) with a Leica DM5500 wide field microscope equipped with a LED Lumencor SOLA lamp, an HCX PL APO 63x Oil objective and an appropriate set of filters (blue, green, orange and far red). We took series of about 40 z-sections with a step of 0.3μm, depending on the thickness of the cell layer in the field.

### Image Processing and Data Analysis

smRNA FISH microscopy images were processed using a custom analysis pipeline based on open source software Ilastik1.3.2 and CellProfiler3.1.8 [65,66]. Briefly, CellProfiler was first used to generate maximum Z-stack projections (all channels) and sum Z-stack projections (Red channel only). Secondly, Ilastik pixel classification projects were used to generate probability maps of dots from max Z-stack projections. One Ilastik workflow has been used for each (Red) and (FarRed) channels using Random Forest Model trained with a mix of images from independent experiments. Finally, CellProfiler was used to segment dots (dots probability maps), cells (sum Z-projections, Red channel), nucleus (max Z-projection, DAPI channel) and cytoplasms (cell minus nucleus), and to relate them with each other. Data analysis was performed using personalised R and Matlab scripts and functions.

### Mathematical models of smFISH data

To model the smFISH average mRNA level we used a least-squares fit of the following function with 2 harmonics for each gene *g*:

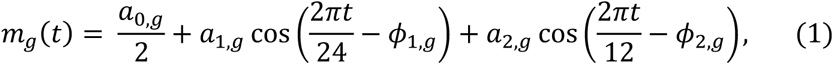

To model the smFISH mRNA count data in a population of cells we used a mixture of negative binomial (NB) distributions. As we explain now, the NB distributions covers the intrinsic noise, while extrinsic noise sources are modelled with mixtures of distributions. The NB is a discrete probability distribution whose parameters can be related to a biophysical model of transcriptional bursting assuming fixed parameters (no extrinsic noise).

For all models, the probability of observing an smFISH count *k* for gene *g* and for cell *i* follows a negative binomial distribution, parameterised with a burst size parameter *b*_*i,g*_ and a bust frequency parameter *f*_*i,g*_

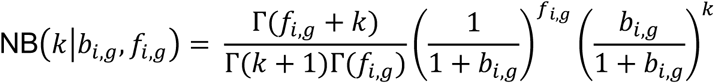

We now describe each the four models M1-M4 (Table 1). In model M1, the burst size (*b*_*g*_) is a constant while the burst frequency (*f*_*g*_) uses the same function *m*_*g*_(*t*) (equation 1) that was fitted to the mean mRNA level but rescaled with a parameter *γ*_*g*_

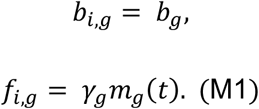

At a given circadian time *t*, all cells share the same parameters. We used weakly informative priors on the parameters *b*_*g*_, *γ*_*g*_ ∼ *N*(0,100).

We next proposed a second model (M2) that incorporates cell size, where we modified the dependence between burst size and cell area with an exponent *β*_*g*_. Consequently, the reformulated model is as follows:

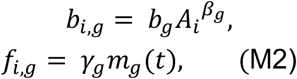

where *A*_*i*_ represents the cell area for cell *I*, and we again used a weakly informative prior *β*_*g*_ ∼ *N*(0,100).

For model M3 we incorporate imperfect synchronisation by allowing each cell to have an individual phase *φ*_*i*_. The amplitude of the oscillatory function is rescaled with the parameter λ

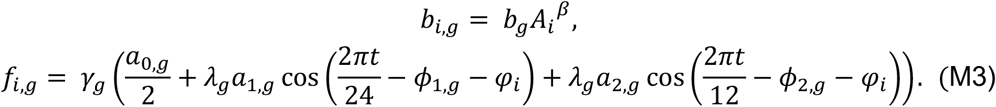

To ensure the cell-specific phases do not become too spread, we used a Von Mises distribution as a prior with a *k* value of 2 and a mean of 0, which has the following density

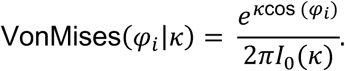

where *I*_*0*_ (*k*) is the modified Bessel function of order 0. We used a uniform prior on λ_*g*_. For our final model (M4) we assume that the burst size in each cell is the product of the cell area and a multiplicative noise term ε_*i,g*_.

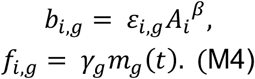

We assume that the noise term ε_*i,g*_ is multivariate log-normally distributed between two genes and is parameterised as follows:

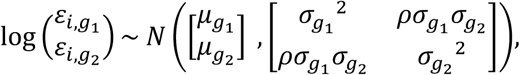

where 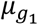 and 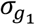 represent the mean and standard deviation for gene 1 and ρ represents the correlation in burst size between genes (in log space). We used an LKJ prior with shape parameter 4 to regularise ρ. The correlation coefficient ρ is inferred between *Bmal1-Cry1* and *Nr1d1-Cry1*, but it is not directly inferred between *Nr1d1* and *Bmal1*. As such, for the 3-dimensional simulations we assume the ε_*i,g*_ parameters are conditionally independent between these two genes (i.e. the entry in the precision matrix for *Nr1d1-Bmal1* is zero).

### Parameter inference and model selection

Having specified the model, we used the Hamiltonian Monte Carlo sampler provided within the STAN probabilistic programming language [37] to sample model parameters from the posterior distribution. For model selection we use PSIS-LOO python function provided with [38].

### Variance decomposition

We decompose the variance by successively conditioning on groups of extrinsic parameters, as shown in Bowsher and Swain [67]. For mRNA counts *Y* and extrinsic parameters *X*_1_, *X*_2_ and *X*_3_, the decomposition is as follows:

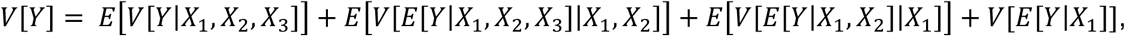

where the first term represents the variance from intrinsic transcriptional noise, and the remaining terms describe the contribution to variance from extrinsic variables. The expectation operator *E* used with conditioning variables (i.e. *E*[*Y*| *X*_*i*_]) denotes averaging over all random variables except those given in the conditioning. In our case the extrinsic conditioning variables *X*_*i*_ correspond to time, cellular area and the burst size noise term ε_*i,g*_. The contribution to variance of each of our extrinsic variables depends on whether the extrinsic conditioning variables *X*_*i*_ is assigned to *X*_1_, *X*_2_ and *X*_3_, and we therefore average over all possible combinations.

The intrinsic transcriptional noise contains the term *V*[*Y*| *X*_1_, *X*_2_, *X*_3_], which is the variance of the negative binomial distribution with fixed values of extrinsic parameters. This term can be further decomposed into a Poisson and a promoter noise component:

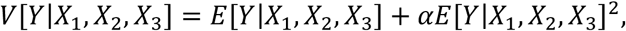

where *α* is the dispersion parameter of the negative binomial. Under our model, the expectation *E*[*Y*| *X*_1_, *X*_2_, *X*_3_] is equal to *bf* and the dispersion parameter *α* is equal to 1/*f*. All terms were computed via simulation of the inferred model, where we used the posterior mean parameter values of the preferred model. All data and code is available at https://github.com/naef-lab/CircadianSMFISH.

## Author Contributions

Conceptualization, N.E.P., A.H., F.N.; Methodology, N.E.P., A.H., J.Y., F.N.; Investigation, A.H., E.D., D.N.; Formal Analysis, N.E.P., A.H.; Data Curation, E.D., A.H.; Software, N.E.P, E.D, A.H, J.Y.; Writing – Original Draft, N.E.P., A.H., F.N.; Writing – Review & Editing, N.E.P., A.H., J.Y., E.D., D.N., F.N.; Resources and supervision, F.N.

## Acknowledgments

We thank the BIOP imaging facility at the EPFL for support in setting up the multichannel smFISH imaging.

