## Supplementary Information for "The circadian oscillator analysed at the single-transcript level"

### The circadian oscillator observed at the single-transcript level - Supplementary Information

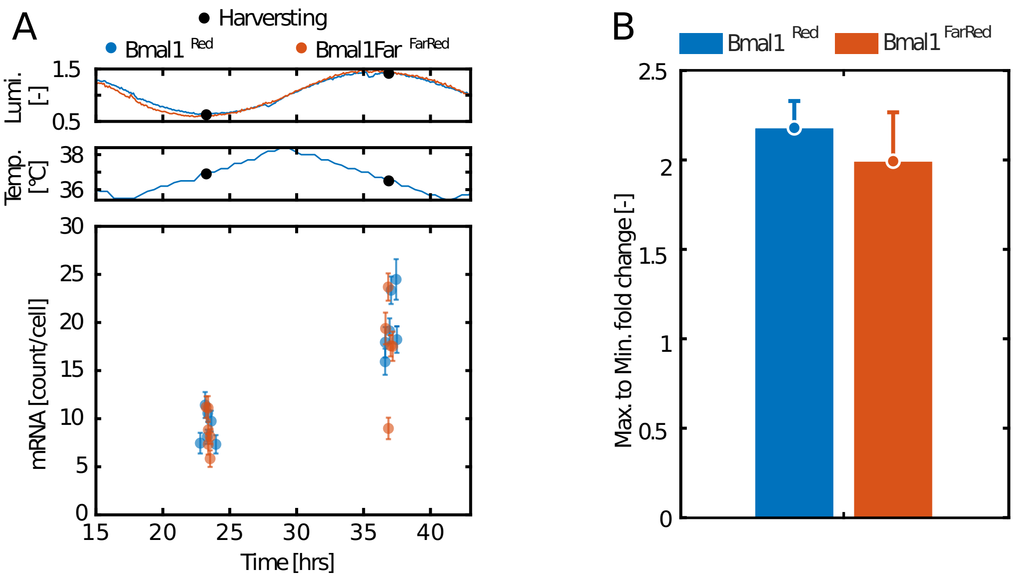


**Supplementary Figure 1:** Amplitude of oscillations with temperature cycle synchronisation. Using a stable Bmal1-sLuc2 NIH3T3 cell line (clone M, Nicolas, 2018), the circadian clock was synchronised using temperature cycles and then split and seeded on a 3.5 cm a Petri dish before recording with an Actimetrics LumiCycle. Note the recording is performed under constant (37°C) temperature. The raw luminescence signal was corrected as described in (Saini et al., 2012). smFISH was performed using probes targeting two non-overlapping regions of *Bmal1* transcripts ($\text{Bmal1}^{\text{Red}}$, $\text{Bmal1}^{\text{FarRed}}$) on three independent cultures of WT NIH-3T3 cells. (A, top) Corrected luminescence from NIH-3T3 cells expressing Bmal1-luciferase. (A, middle) Temperature profile recorded by the Memert incubator. (A, bottom) Mean mRNA count for each replicate. Error bars represent the standard error. (B) Peak-to-trough fold change. Error bars are standard deviation.

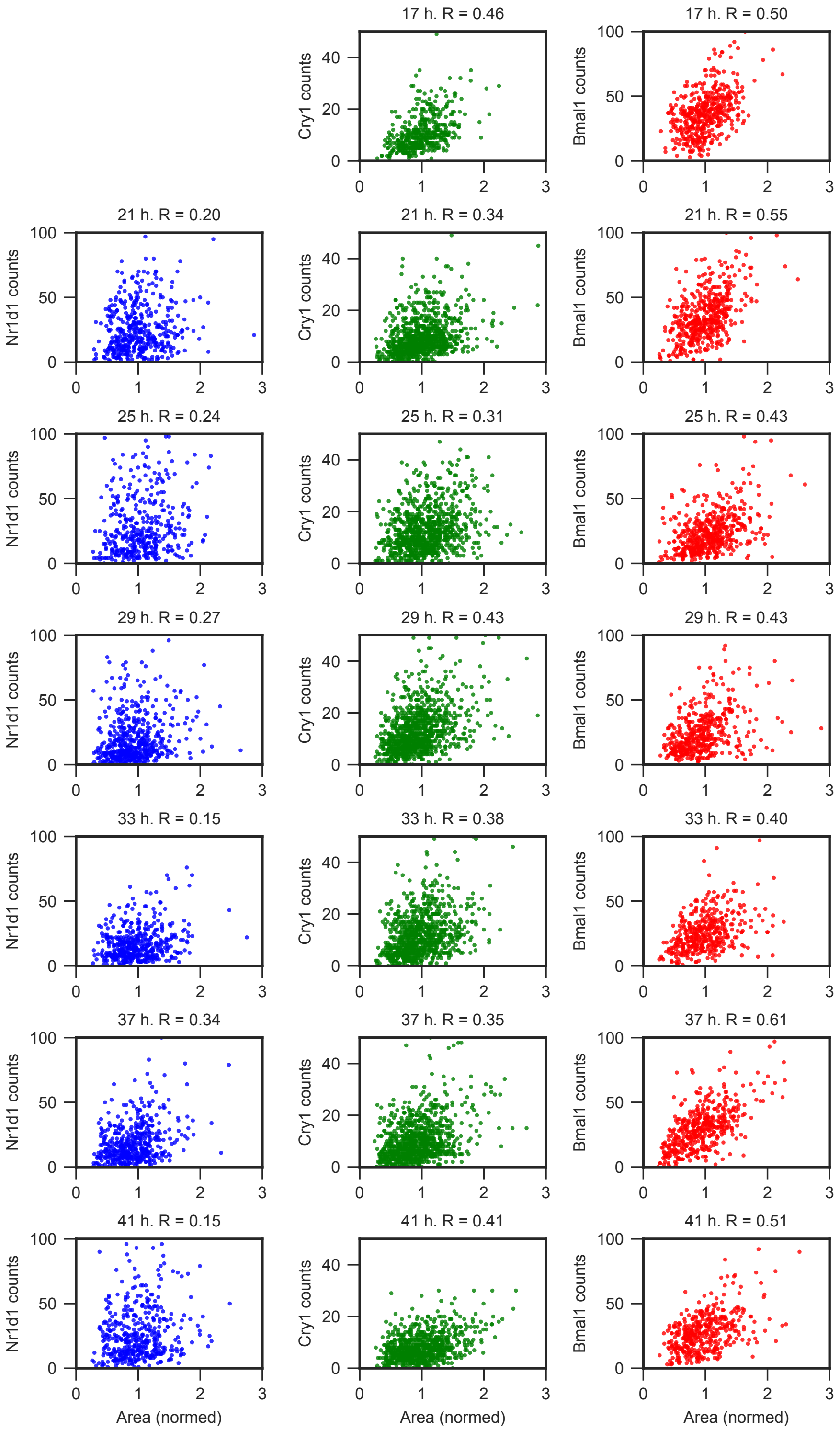


**Supplementary Figure 2:** mRNA count number as a function of area for *Nr1d1*, *Cry1* and *Bmal1* at each time point. The area is normalised such that the mean is 1.

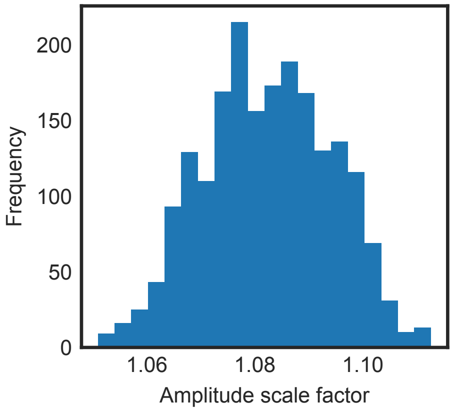


**Supplementary Figure 3:** Posterior parameter estimates for the amplitude scale factor $\lambda$ in model M3 (described in Methods). A scale factor of 1.08 corresponds to a 8% increase in amplitude relative to model M1.

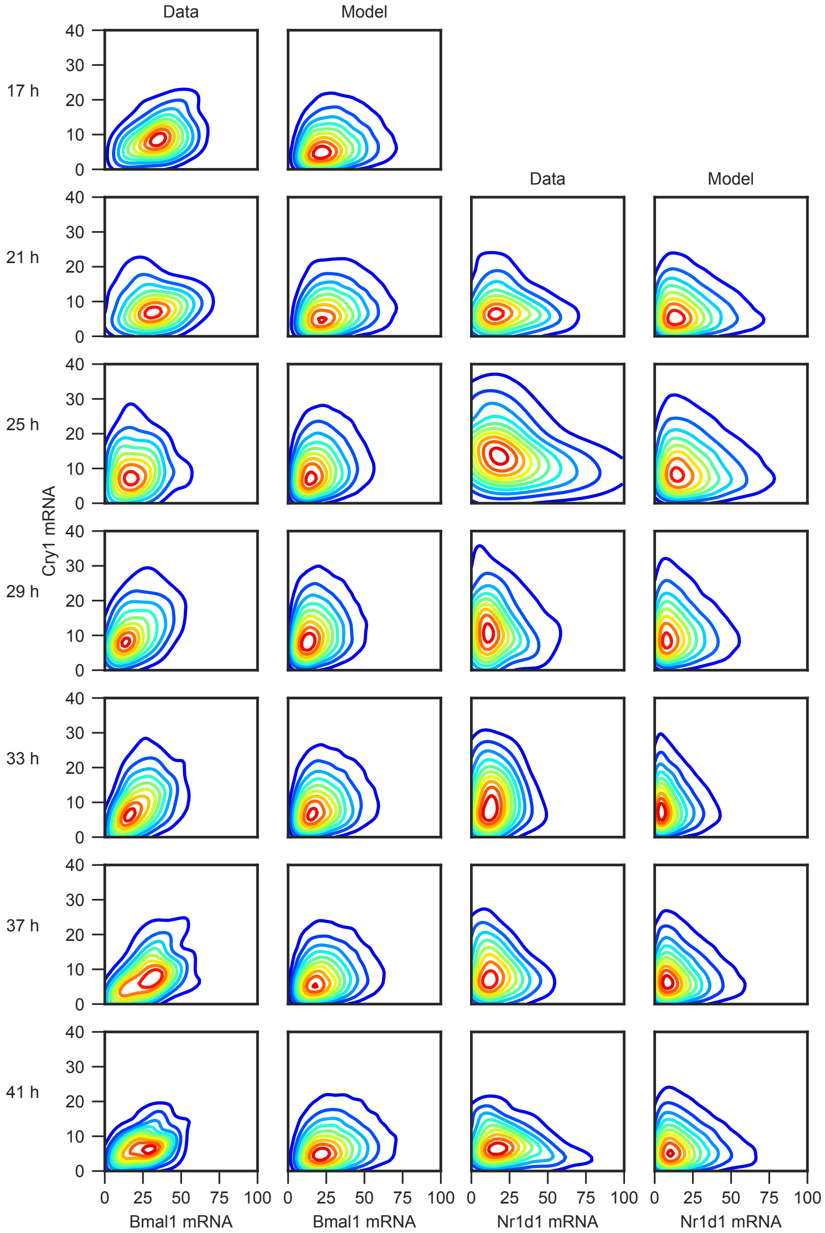


**Supplementary Figure 4:** Comparison of the kernel density estimate of the data with model M4 for all time points. To estimate the probability from the model, the model was simulated 15 times using the posterior mean parameter values combined with the measured cell areas.
